# A synthetic bispecific antibody capable of neutralizing SARS-CoV-2 Delta and Omicron

**DOI:** 10.1101/2022.01.04.474803

**Authors:** Tom Z. Yuan, Carolina Lucas, Valter S. Monteiro, Akiko Iwasaki, Marisa L. Yang, Hector F. Nepita, Ana G. Lujan Hernandez, Joseph M. Taft, Lester Frei, Sai T. Reddy, Cédric R. Weber, Kevin P. Malobisky, Rodrigo Mesquita, Aaron K. Sato

**Author notes:** To whom correspondence should be addressed: Aaron Sato.

## Abstract

Bispecific antibodies have emerged as a promising strategy for curtailing severe acute respiratory syndrome coronavirus 2 (SARS-CoV-2) immune escape. This brief report highlights RBT-0813 (also known as TB493-04), a synthetic, humanized, receptor-binding domain (RBD)-targeted bispecific antibody that retains picomolar affinity to the Spike (S) trimers of all major variants of concern and neutralizes both SARS-CoV-2 Delta and Omicron *in vitro*.

## Introduction

Since its emergence in December 2019, SARS-CoV-2 continues to evolve substantially, acquiring sets of mutations that enhance the virus’s potency, transmissibility, infectivity, and ability to escape natural and acquired immunity.^1^ Virtually all of these fitness-enhancing mutations are found in Spike (S), the protein through which SARS-CoV-2 binds to angiotensin-converting enzyme 2 (ACE2) on the host cell surface during infection. The receptor-binding domain (RBD) of S is the primary target of neutralizing antibodies produced after natural infection or vaccination against SARS-CoV-2; this fact has fuelled speculation that SARS-CoV-2 could escape natural and acquired immunity through mutations in the S.^2^ Several variants of concern possessing mutations in the RBD have displayed varying degrees of immune escape, including the Beta (B.1.351), Gamma (P.1) and Delta (B.1.617.2) variants.^3–6^ The recently detected Omicron (B.1.1.529) variant, which possesses at least 30 amino acid mutations in S alone,^7^ is particularly concerning, as early reports show clear reductions in the efficacy of current therapeutics, including those monoclonal antibody cocktails with Emergency Use Authorization, as well as the commonly used two-dose mRNA vaccine regimens.^8–21^ SARS-CoV-2 escape mutations in the RBD presents a particular risk for recently developed neutralizing antibody therapeutics and endangers ongoing public health responses, thus underscoring the need for new therapeutic approaches, such as multivalent antibodies, which have recently been shown to potentiate SARS-CoV-2 neutralization and reduce immune escape when compared to monovalent antibodies.^22–26^ Because the Delta and Omicron variants account for 99.7% of cases in the United States (as of December 25, 2021^27^), a bispecific antibody capable of binding and neutralizing both variants would represent a therapeutic and public health advantage over currently available treatment modalities.

In this paper, we highlight RBT-0813, a synthetic VHH bispecific antibody capable of binding and neutralizing the SARS-CoV-2 Delta and Omicron variants. This bispecific antibody links together two humanized VHH antibodies — TB202-03 and TB339-031 — with constant heavy chain 2 (CH2) and 3 (CH3) Fc domains (**Figure 1**). TB202-03, which was discovered by panning the TB202 VHH library^28^ against the S1 monomer of the SARS-CoV-2 WA1 strain, has been shown to effectively neutralize pseudoviruses encoding the Alpha, Beta, and Gamma S proteins, but not those encoding L452R-bearing S protein variants such as Delta and Epsilon (B.1.429). TB339-031 was discovered by panning the TB201 VHH library^28^ against the Beta S1 (subdomain of S) and was found to bind and neutralize L452R-bearing S protein variants. Here, we describe the biophysical and functional characterization of the bispecific RBT-0813 antibody, focusing on its binding and neutralization of Delta and Omicron variants.

**Figure 1.**
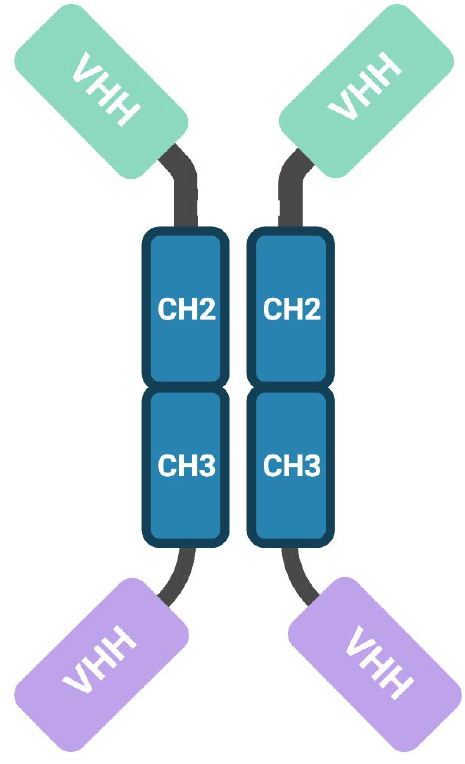
Schematic of RBT-0813, a synthetic bispecific antibody. Green, TB202-03; purple, TB339-031; blue, Fc domains.

## Results

### RBT-0813 binds SARS-CoV-2 Delta and Omicron with picomolar affinity

RBT-0813 was constructed from lead VHH antibodies discovered in biopanning campaigns against the ancestral (TB202-03) and Beta (TB339-031) S proteins. To create a broadly neutralizing antibody, we combined leads TB202-03 and TB339-031 into a single bispecific construct. Screening by surface plasmon resonance (SPR) revealed picomolar apparent binding affinities between RBT-0813 and the prefusion-stabilized S trimers of the Alpha, Beta, Delta, Gamma, Kappa (B.1.617.1), and Omicron variants (**Figure 2a, Table 1**). SPR traces obtained with TB202-03 and TB339-031 showed which of the two contributed to RBT-0813’s binding to each S trimer. Although TB202-03 bound to the Alpha, Beta, Gamma, Kappa with the same affinity as the ancestral S trimer, it displayed reduced (yet still picomolar) affinities with the Kappa and Delta variants (**Figure 2a, Table 1**). By contrast, TB339-031 bound every S trimer variant with low picomolar affinities except the Omicron variant (**Figure 2a, Table 1**). Fortuitously, TB202-03 displayed low picomolar affinity to the S trimer of Omicron, as did RBT-0813. SPR experiments performed using variant S1 monomers showed the same patterns, although apparent binding affinities were in the nanomolar range (**Figure 2a**). These data agree with previous biophysical and functional characterizations of TB202-03^28^ (CoVIC-094 in Hastie et al.^29^) that showed reduced activity of TB202-03 against S proteins bearing the L452R mutation (namely, Delta and Epsilon).

**Figure 2.**
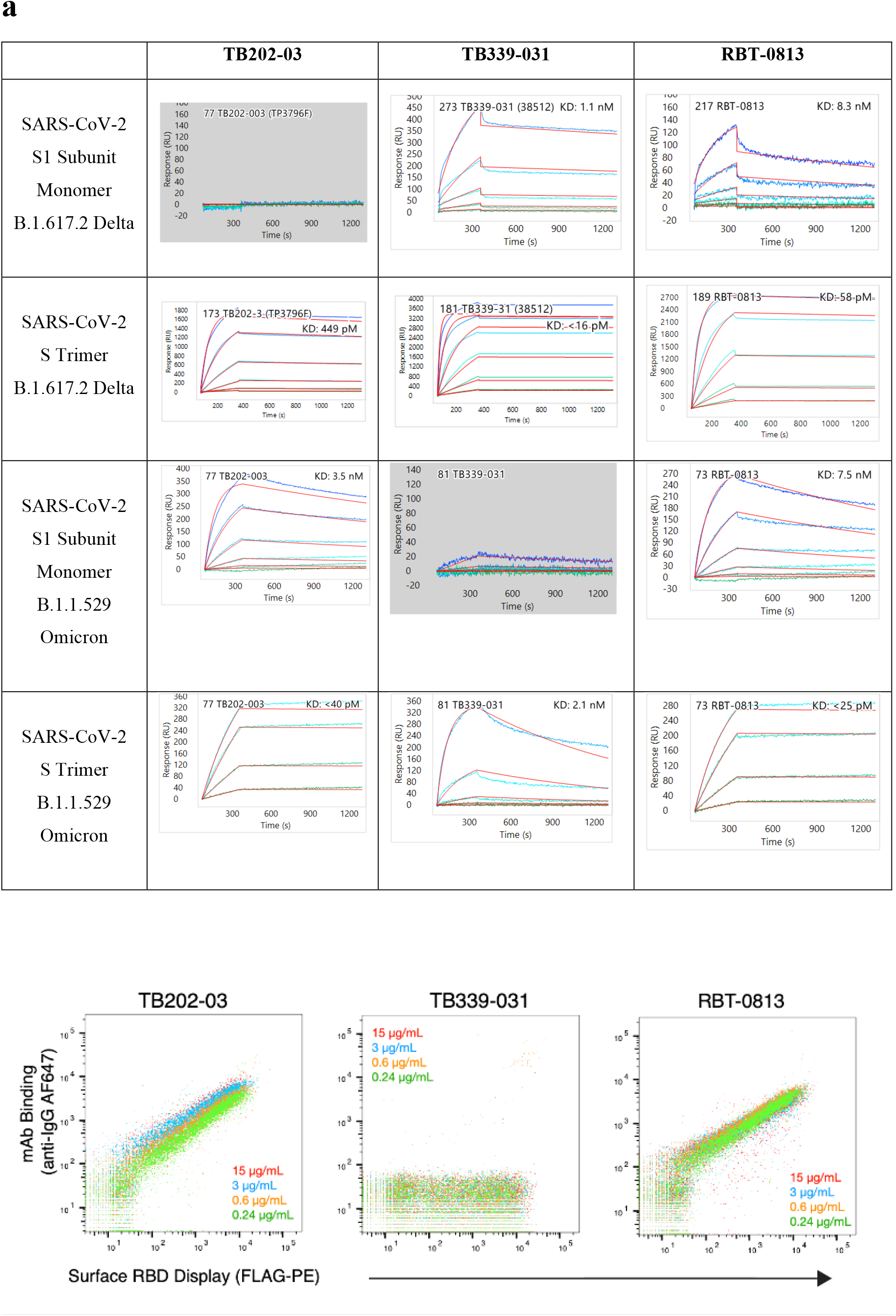
Biophysical characterization of TB493-04 (RBT-0813) and its parental constructs TB202-03 and TB339-031. (a) SPR traces of TB202-03, TB339-031, and RBT-0813 binding to the S1 monomers and S trimers of SARS-CoV-2 Delta and Omicron. (b) Binding of RBT-0813 to Omicron S1 RBD displayed on the surface of yeast. Binding of Omicron S1 RBD by RBT-0813 is confirmed by high fluorescence in both channels.

**Table 1.**
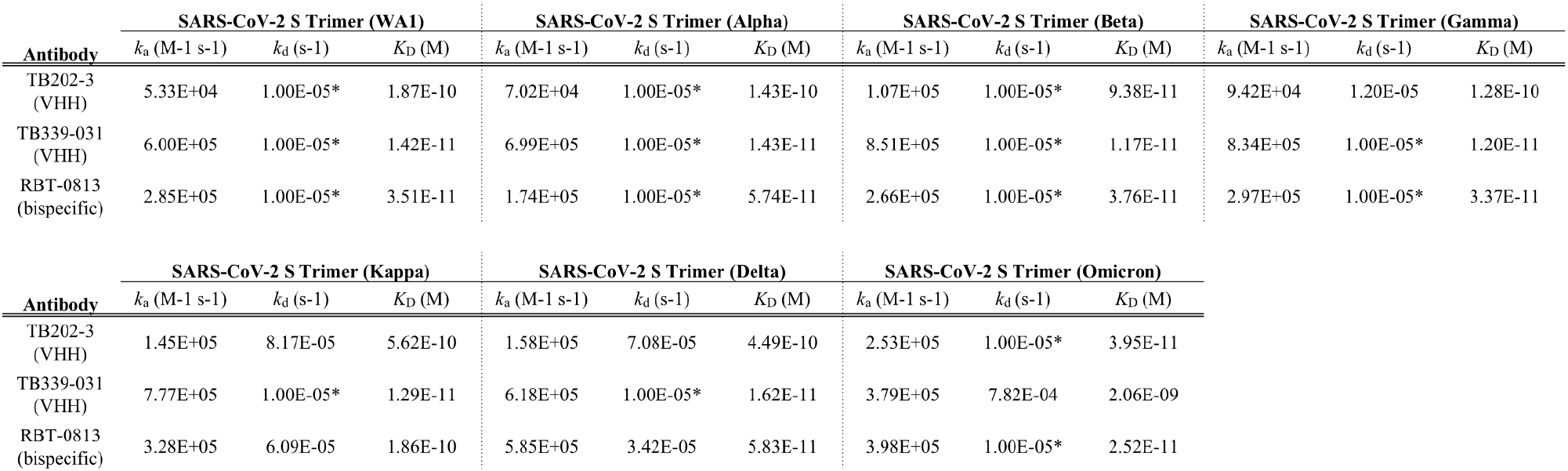
Apparent binding affinity summary of TB493-04 (RBT-0813) and its parental constructs TB202-03 and TB339-031 to S glycoprotein prefusion trimers as measured by SPR. Data were analyzed in Carterra’s Kinetics Tool software with 1:1 binding model. *Indicates off-rate is slower than limit of detection.

Consistent with the SPR data described above, TB202-03 and RBT-0813, but not TB339-031, bound the Omicron S1 RBD displayed on the surface of yeast, as measured by flow cytometry (**Figure 2b**); thus providing independent confirmation that TB202-03 mediates the Omicron binding of RBT-0813.

### RBT-0813 potently neutralizes SARS-CoV-2 Delta and Omicron

According to early reports, the vast majority of monoclonal antibodies in development fail to neutralize Omicron, including antibodies that effectively neutralize Delta.^9,18,19^ Similarly, the U.S. Food and Drug Administration (FDA) has recently updated the fact sheets for health care providers for the EUA of antibody therapies from Eli Lilly^33^ (cocktail of two monoclonal antibodies: bamlanivimab and etesevimab) and Regeneron Pharmaceuticals^34^ (cocktail of two monoclonal antibodies: casirivimab and imdevimab). In both of these cases, in vitro binding and neutralization studies have suggested that neither of the antibody cocktail therapies are likely to have clinical efficacy against the Omicron variant. To assess the neutralization potential of RBT-0813, we utilized authentic viruses isolated from nasopharyngeal specimens of patients in plaque reduction neutralization assays to determine whether RBT-0813 can neutralize SARS-CoV-2 Ancestral, Delta and Omicron. As shown in **Figure 3**, RBT-0813 neutralizes authentic Delta and Omicron at half maximal effective concentrations (EC50) of 521.1 and 713.6 ng/ml, respectively. In line with the binding data, RBT-0813 neutralizes Delta and Omicron through either the TB339-031 and TB202-03 VHH antibodies, respectively. These values are comparable to those obtained with sotrovimab (VIR-7831), one of the few monoclonal antibodies that shows effective neutralization against both Delta and Omicron at 325 and 917 ng/ml, respectively.^18^ In a separate study, sotrovimab neutralized Ancestral and Omicron at 179 and 320 ng/ml, respectively.^9^ Interestingly, RBT-0813 neutralizes Ancestral SARS-CoV-2 at EC50 of 104.4 ng/mL, which is approximately 5-fold and 7-fold enhanced compared to Delta and Omicron, respectively. Since TB202-03 and TB339-031 both bind Ancestral SARS-CoV-2 with high affinity, they avidly work together to neutralize in concert with one another.

**Figure 3.**
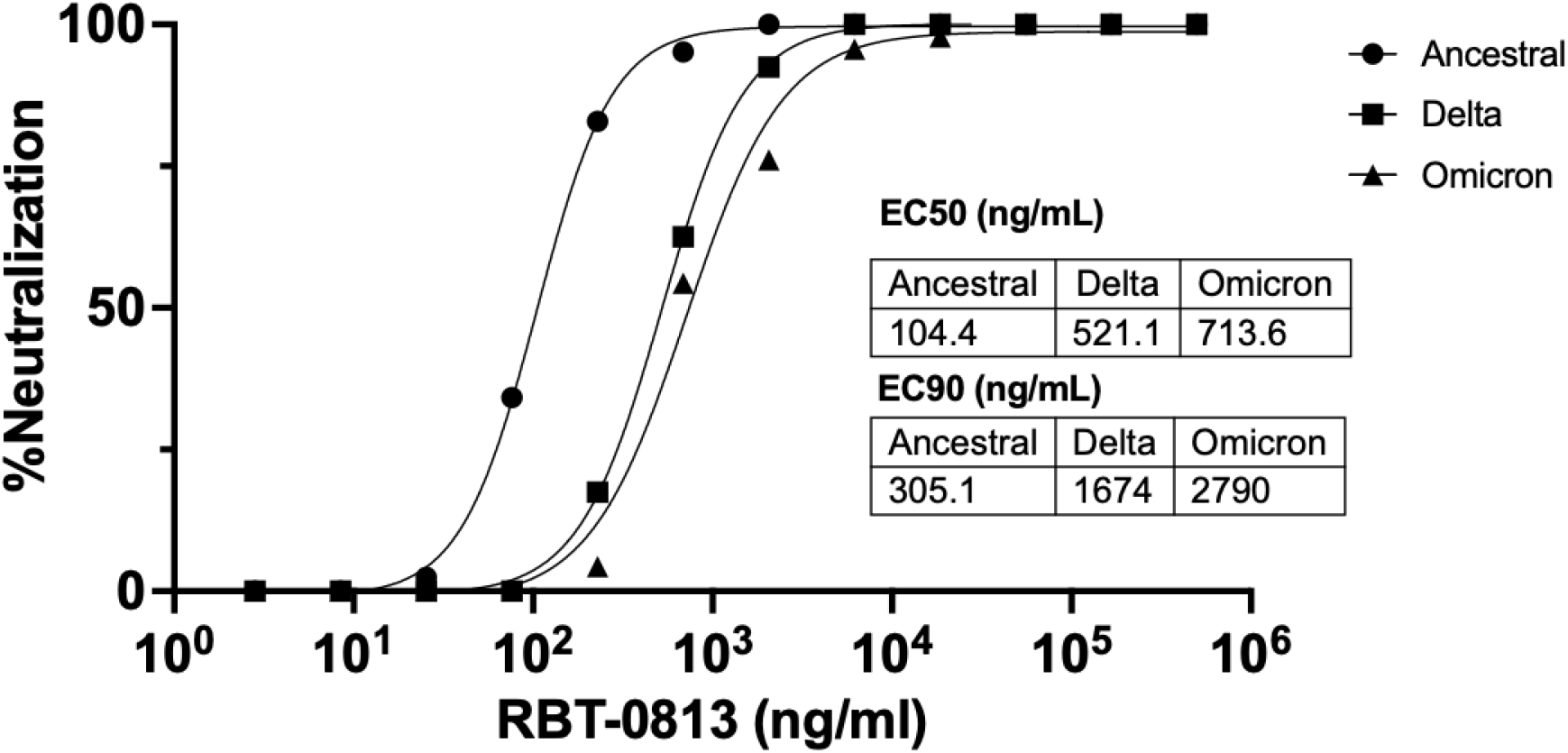
Neutralization of authentic SARS-CoV-2 Delta and Omicron by RBT-0813. Neutralization curves against ancestral lineage A virus (WA1,USA), Delta and Omicron SARS-CoV-2 variants. Antibody-mediated neutralization was accessed by SARS-CoV-2 infection on TMPRSS2-VeroE6 cells. EC50 and EC90 values represent the concentrations required to reduce the number of plaques by 50% and 90%, respectively. Representative curves of three independent experiments with technical duplicates.

## Discussion

First detected in November 2021, the Omicron variant has quickly spread worldwide, causing infections in at least 89 countries.^30^ Omicron’s rapid transmission and sheer number of S mutations, especially in the RBD, quickly sparked concerns about the variant and its ability to escape immune protection and existing antibody therapeutics. Moreover, despite Omicron’s meteoric rise, with its displacement of Delta in some areas, Delta remains a major threat to public health, especially because early animal studies indicate that Delta causes more severe disease than Omicron.^31^ In this report, we show that RBT-0813 binds and neutralizes not only Delta, which has only recently begun to lose its foothold as the dominant global strain, but importantly, it binds and neutralizes Omicron as well. With this report, RBT-0813 represents one of the few reported Abs that retain significant antiviral activity against Omicron.

Most of the mAbs that have been shown to neutralize Omicron target conserved *Sarbecovirus* epitopes and do not compete with ACE2 for S binding.^9,18,19^ Although the mechanism through which RBT-0813 neutralizes Omicron has not yet been clarified, the TB202-03 arm presumably contributes more than the TB339-031 arm due to its higher affinity to the Omicron S1 monomer and S trimer (**Figure 2a**). In stark contrast to antibodies that target a highly conserved *Sarbecovirus* epitope (e.g., sotrovimab), TB202-03 binds toward the outer edge of the receptor-binding motif of SARS-CoV-2 (the RBD-4 community in Hastie et al.^29^), does not bind SARS-CoV S1, and competes with ACE2 for SARS-CoV-2 S binding.^28^ The antiviral activity of RBT-0813 may be explained by the fact that none of the key residues in the TB202-03 epitope — namely, N450, I472, and F490^28^ — are mutated in Omicron.

In aggregate, these early data highlight RBT-0813 as a promising, innovative bispecific VHH therapeutic candidate for further development in the ongoing SARS-CoV-2 pandemic.

## Materials and Methods

### SPR affinity measurements

SPR experiments were performed on a Carterra LSA SPR biosensor equipped with a HC30M chip at 25°C in HBS-TE. Antibodies were diluted to 10 μg/mL and amine-coupled to the sensor chip by EDC/NHS activation, followed by ethanolamine HCl quenching. Increasing concentrations of analyte were flowed over the sensor chip in HBS-TE with 0.5 mg/mL BSA with 5 minute association and 15 minute dissociation. SARS-CoV-2 protein reagents were sourced commercially from Acro Biosystems: S1 B.1.1.529 Omicron (S1N-C52Ha), S Trimer B.1.1.529 Omicron (SPN-C52Hz), S1 B.1.617.2 Delta (S1N-C52Hu), S Trimer B.1.617.2 Delta (SPN-C52He), S Trimer WA1 (Acro SPN-C52H9), S Trimer B.1.1.7 Alpha (Acro SPN-C52H6), S Trimer B.1.351 Beta (Acro SPN-C52Hk), S Trimer P.1 Gamma (Acro SPN-C52Hg) and S Trimer B.1.617.1 Kappa (Acro SPN-C52Hr). Following each injection cycle, the surface was regenerated with 2× 30-second injections of IgG elution buffer (Thermo). Data were analyzed in Carterra’s Kinetics Tool software with 1:1 binding model.

### Flow binding assay

#### Growth/induction

EBY100 yeast cells transformed with pYD1-Omicron RBD (amino acids 331-531) were picked from a selective plate and inoculated in 1 mL SD-UT medium [yeast nitrogen base-casamino acids (YNB-CAA) (BD Biosciences 223120) + 2% D-(+)-Glucose (Sigma G5767-500G) growth medium including phosphate buffer (5.4 g/l Na_2_HPO_4_, 8.6 g/l NaH_2_PO_4_·H_2_O) and 1X Penicillin-Streptomycin]. The cultures were incubated for 12-18 h at 30 °C and 250 RPM. 2 x 10^7^ cells were pelleted by centrifugation for 30 s at 8,000 x g. The pellets were resuspended in 1 mL SG-UT [(YNB-CAA) (BD Biosciences 223120) + 2% D-(+)-Galactose (Sigma G0625-500G) induction medium including phosphate buffer (5.4 g/l Na2HPO4, 8.6 g/l NaH2PO4·H2O) and 1X Penicillin-Streptomycin]. The induction cultures were incubated for 32-40 h at 23 °C and 250 RPM.

#### Staining

6 x 10^5^ cells were added into the wells of a 96 well filter plate (MSHVS4510 Millipore MultiScreenHTS HV Filter Plate, 0.45 μm, clear, sterile). Each of the 3 constructs (TB202-03, TB339-031, and RBT-0813) were diluted to final concentrations with DPBS + 0.5% BSA + 2mM EDTA + 0.1% Tween20. 20 μL of mAb solution was added to the cells in the filter wells and the cells were incubated for 1 h at 4 °C and 750 RPM on a thermomixer (Eppendorf, ThermoMixer C). After the incubation, the liquid in the plate was removed using a vacuum manifold. The pressure was kept at <5 bar. To wash the cells, 200 μL DPBS + 0.5% BSA + 2mM EDTA + 0.1% Tween20 was added to each well and subsequently removed using the vacuum manifold.

20 μL anti-IgG AlexaFluorophore 647 mAb (5 μg /mL) (Jackson Immunoresearch 109-605-098) was added to the cells and they were incubated for 45 min at 4 °C and 750 RPM on a thermomixer (Eppendorf, ThermoMixer C). The liquid in the plate was removed using a vacuum manifold, and the wash step was repeated.

For the expression staining, 20 μL anti-FLAG PE mAb (1 μg/mL) (Biolegend 637309) was added to the cells and they were incubated for 30 min at 4 °C and 750 RPM on a thermomixer (Eppendorf, ThermoMixer C). The liquid in the plate was removed using a vacuum manifold. No wash step was performed. 200 μL DPBS + 0.5% BSA + 2mM EDTA + 0.1% Tween20 was added to each well and the cells were resuspended by repeated pipetting. 20 μL of resuspended cells were mixed with 180 μL DPBS + 0.5% BSA + 2mM EDTA + 0.1% Tween20.

#### Scanning

The cells were scanned using a BD Fortessa analyzer equipped with an HTS system. The following lasers and bandpass (BP) filters were used: 561 nm with a BP filter of 586/15 and 640 nm with a BP filter of 670/14. For each sample, 10^4^ events were measured. The data was analyzed using FlowJo V10.4.2.

### SARS-CoV-2 viral culture

TMPRSS2-VeroE6 kidney epithelial cells were cultured in Dulbecco’s Modified Eagle Medium (DMEM) supplemented with 1% sodium pyruvate (NEAA) and 10% fetal bovine serum (FBS) at 37°C and 5% CO2. The cell line has been tested negative for contamination with mycoplasma. SARS-CoV-2 ancestral strain, lineage A(USA-WA1/2020), was obtained from BEI Resources (#NR-52281). Delta and Omicron variants were isolated from nasopharyngeal specimens as previously described^32^. Expanded viruses were re-sequenced following the same method as described above and genome sequences were uploaded to GenBank and the aligned consensus genomes are available on GitHub (https://github.com/grubaughlab/paper_2021_Nab-variants). The pelleted virus was then resuspended in PBS and aliquoted for storage at −80°C. Viral titers were measured by standard plaque assay using TMPRSS2-VeroE6. All experiments were performed in a biosafety level 3 laboratory with approval from the Yale Environmental Health and Safety office.

### Authentic virus neutralization assay

Serial dilutions of RBT-0813 (500 μg/ml to 2.89 ng/ml) were individually incubated with the ancestral SARS-CoV-2 strain (USA-WA1/2020), the Delta variant, or Omicron variant, for 1 h at 37 °C. (Viral concentrations were optimized to generate 60-120 plaques per well.) The resulting mixtures were then applied to TMPRSS2-VeroE6 cells, plated in a 12-well plate, for 1 hr, after which MEM supplemented with NaHCO_3_, 4% FBS, and 0.6% Avicel, was added to each well. At 40 h post-infection, cells were fixed with 10% formaldehyde for 1 h and then stained with 0.5% crystal violet to resolve plaques. All experiments were performed in parallel with baseline controls. Analyses of plaque counts were done using GraphPad Prism software, version 8.4.

## Competing Interests

Twist Biopharma paid Alloy Therapeutics/ Department of Biosystems Science and Engineering, ETH Zurich to generate referenced data.

A.S and T.Y. are paid employees and stockholders of Twist Bioscience Corporation that has licensed RBT 0813 for a substantial equity interest in Revelar Biotherapeutics, Inc. and other contingent consideration.

K.M. and R.M. are paid employees and stockholders of Revelar Biotherapeutics., Inc.

A.I. serves as the Co-Chair of Revelar Biotherapeutics, Inc.’s Scientific Advisory Board and is a stockholder.

The other authors declare no competing interests.

